# Reanalysis of single-cell RNA sequencing data does not support herpes simplex virus 1 latency in non-neuronal ganglionic cells in mice

**DOI:** 10.1101/2023.07.17.549345

**Authors:** Werner J.D. Ouwendijk, Pavitra Roychoudhury, Anthony L. Cunningham, Keith R. Jerome, David M. Koelle, Paul R. Kinchington, Ian Mohr, Angus C. Wilson, Georges M.G.M. Verjans, Daniel P. Depledge

## Abstract

Most individuals are latently infected with herpes simplex virus type 1 (HSV-1) and it is well-established that HSV-1 establishes latency in sensory neurons of peripheral ganglia. However, it was recently proposed that latent virus is also present in immune cells recovered from ganglia in a mouse model used for studying latency. Here, we reanalyzed the single-cell RNA sequencing (scRNA-Seq) data that formed the basis for this conclusion. Unexpectedly, off-target priming in 3’ scRNA-Seq experiments enabled the detection of non-polyadenylated HSV-1 *latency-associated transcript* (*LAT*) intronic RNAs. However, *LAT* reads were near-exclusively detected in a mixed population of cells undergoing cell death. Specific loss of HSV-1 *LAT* and neuronal transcripts during quality control filtering indicated widespread destruction of neurons, supporting the presence of contaminating cell-free RNA in other cells following tissue processing. In conclusion, the reported detection of latent HSV-1 in non-neuronal cells is best explained by inaccuracies in the data analyses.

## Introduction

All herpesviruses share the ability to establish a lifelong latent infection in their hosts, which facilitates virus reactivation and intermittent spread to naïve hosts. Herpesviruses belonging to the *Alphaherpesvirinae* subfamily, which includes the ubiquitous human pathogen herpes simplex virus type 1 (HSV-1), establish latency in neurons of the peripheral nervous system [1]. The primary site of HSV-1 latency are sensory neurons of the trigeminal ganglia (TG) and/or dorsal root ganglia (DRG) [2]. Moreover, HSV-1 DNA also persists in neurons of other sensory and autonomic ganglia [3,4], and the possible existence of a latent HSV-1 reservoir in the cornea has been a long-standing debate in the field [5,6]. Therefore, the development of single-cell RNA sequencing (scRNA-Seq) technologies provides an unique opportunity to study latency and reactivation in neuronal latency models [7], as well as to provide evidence for whether HSV latency is established in non-neuronal cells. A recent study by Wang *et al*. [8] addressed the latter and concluded that, in addition to neurons, HSV-1 establishes latency in immune cells that are present in TG of HSV-1 experimentally-infected mice. Here, we present a reanalysis of the scRNA-Seq data from Wang *et al*. [8] that demonstrates inaccuracies in their data analyses and that argue against this conclusion.

During latency, HSV-1 gene expression is highly restricted and limited to the *latency-associated transcript* (*LAT*) and associated miRNAs [9–14]. The primary *LAT* transcript is 8.3 kb in size, capped, and polyadenylated [15–18]. Splicing produces stable 1.5 and 2 kb *LAT* intron lariats that accumulate to high levels in sensory neurons, while the highly unstable 6.3 kb spliced polyadenylated RNA is rapidly processed into viral microRNAs [19,20] (for a comprehensive review of *LAT*, see [21]). Two major claims are reported by Wang *et al*: (i) in addition to neurons, various types of immune cells recovered from TG of experimentally infected mice express HSV-1 *LAT*, and (ii) the presence of *LAT* in these cells indicates that HSV-1 can establish latency in non-neuronal cells present in the TG. The core data supporting these claims was obtained by droplet-based scRNA-Seq analysis (10X Genomics platform) of TG from uninfected C57BL/6 mice (dataset: “Uninf-1”) and two biological replicate groups of C57BL/6 mice infected via the corneal route with 2 × 10^5^ pfu/eye of HSV-1 strain McKrae 35 days earlier (datasets: “Inf-1” and “Inf-2”). Each biological replicate was obtained by pooling the paired left and right TG from 15 animals (i.e., 30 TG per biological replicate). To examine the claims in more detail, we aimed to reproduce the analyses presented by Wang *et al*. [8]. Although the neither the (raw) scRNA-Seq datasets nor the analyses scripts were available upon publication or currently linked to the online article, we obtained the raw data from the study (SRA PRJNA937697, GEO GSE225839) via the handling editor. What follows is a reanalysis of the data presented by Wang *et al*. [8] using the same filtered barcode matrices that serve as input for the scRNA-Seq analysis.

## Results

### Quality control of scRNA-Seq datasets

Isolation of dissociated, single cells from organs requires mechanical and/or enzymatic tissue dissociation, typically followed by removal of dead cells and (if needed) further purification of cells of interest by magnetic bead-or flow cytometry-based cell sorting. Quality control (QC) filtering of the obtained scRNA-Seq datasets is therefore a critical first step [22]. The Chromium Single Cell 3’ v3.1 Reagent Kit (10X Genomics) used by Wang *et al*. [8] for library preparation is designed to capture polyadenylated RNAs and prime reverse transcription using a poly(T) primer that also includes the barcode and unique molecular index (UMI) sequences. QC filtering involves the identification and removal of doublets, as well as an assessment of cell viability in each of the samples. This latter is achieved by measuring, for each individual cell (i) the number of unique genes detected, (ii) the total number of RNA molecules (UMI) recovered and (iii) the proportion of reads derived from mitochondrial RNAs [23,24] (**Fig. 1**). In the original matrix count files generated by Wang *et al*. [8], the datasets designated Inf-1 and Uninf-1 displayed relative similar results with median counts of 1,979 and 1,822 distinct genes detected per cell, a median total RNA count of 7,148 and 6,879, and with 51% and 83% cells having a mitochondrial RNA fractions count < 15%. By contrast, dataset Inf-2 showed very different results with a median of only 558 distinct genes per cell, a median total RNA count of just 929, and only 33% of cells having a mitochondrial RNA fraction < 15% (**Fig. 1A**). These data indicate a high proportion of dying/dead cells within the Inf-2 dataset. Next, we applied filters on mitochondrial RNA content and unique gene counts, according to the parameters described by Wang *et al*. [8], namely that cells were only retained if between 300 – 9,000 distinct genes were detected, and the proportion of mitochondrial reads present was below 15%. At this stage, we observed large numbers of low-quality cells filtered out of each dataset (**Fig. 1B**). This resulted in 3,608 cells for Uninf-1 (reduced from 4,206, a loss of 14%), 3,158 cells for Inf-1 (reduced from 6,155, a loss of 49%), and 5,660 cells for Inf-2 (reduced from 17,014, a loss of 67%).

**Figure 1.**
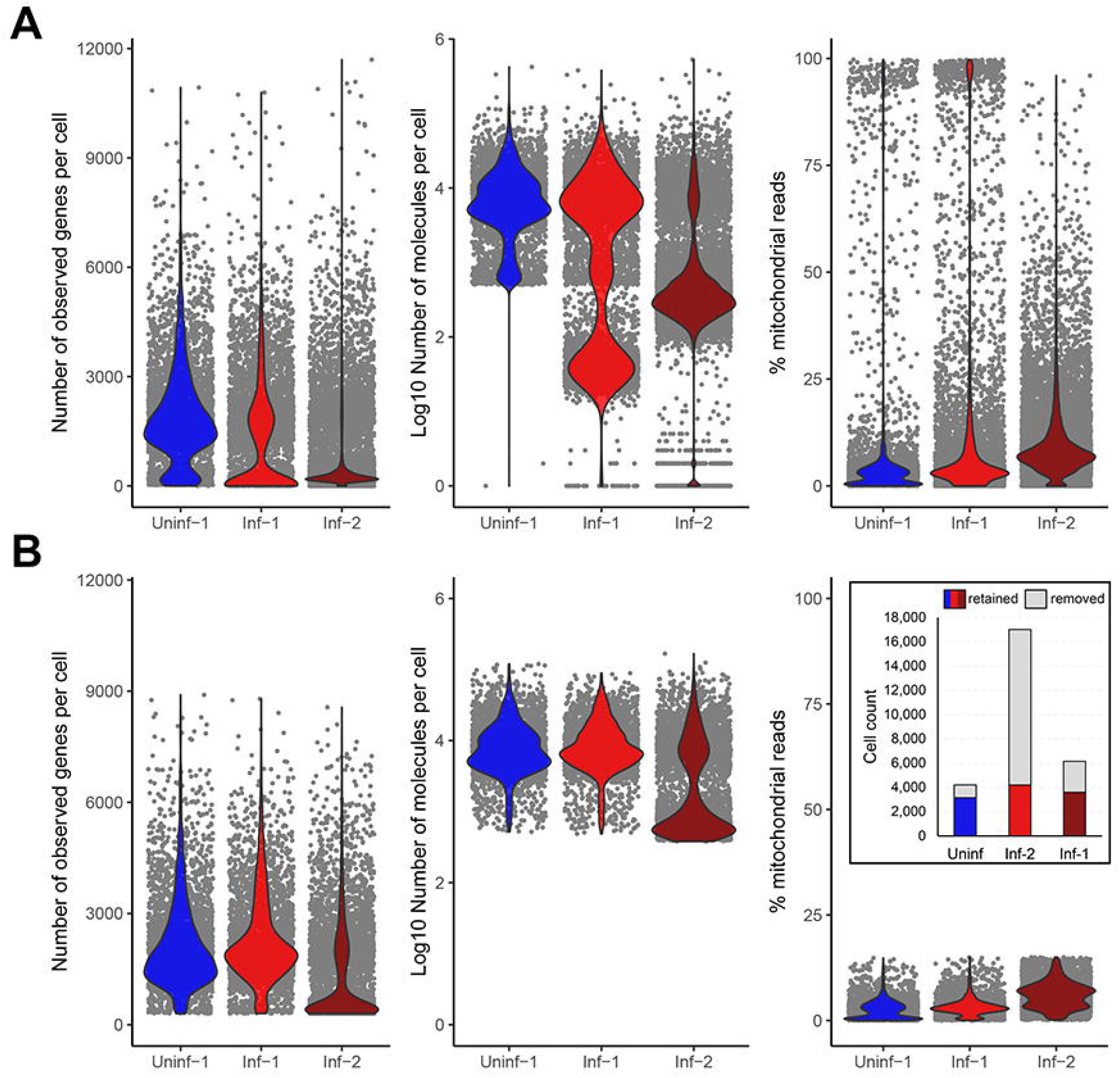
Quality control of scRNA-Seq datasets derived from pools of HSV-1 latently-infected mouse trigeminal ganglia. **(A)** Using the filtered barcode matrices generated by Wang *et al*. [8], the quality of each dataset was assessed by (left) the number of unique genes detected per cell, (middle) the total number of RNA molecules (UMI) recovered per cell and (right) the proportion of reads per cell derived from mitochondrial RNAs. **(B)** Quality control filtering of these datasets dramatically reduced the total number of cells available for analysis, indicative that many dead/dying cells were present in the original single-cell suspensions of Inf-1 and Inf-2. Filtering parameters removed cells with less than 300 or more than 9000 distinct expressed genes, and cells for which more than 15% of reads derived from mitochondrial RNAs. Inset: Number of cells before and after filtering.

### Clustering and annotation of single cells

One of the most contentious components of any scRNA-Seq analyses is the reduction of multi-dimensional into simple two-dimensional figures through either t-distributed stochastic neighbor embedding (t-SNE) or Uniform Manifold Approximation and Projection (UMAP) techniques. While this has been reviewed elsewhere [22] it is worth noting that significant care must be taken when trying to interpret these data. Here, integration of the three datasets and clustering was performed in a similar manner to Wang *et al*. [8], matching as many parameters as possible (see Methods) (**Fig. 2A**). A deeper analysis of the clusters revealed significant differences in the relative proportions of cell types present in each dataset with some clusters being almost entirely derived from a single dataset (e.g., Cluster 0 and to a lesser extent cluster 2, **Fig. 2B**).

**Figure 2.**
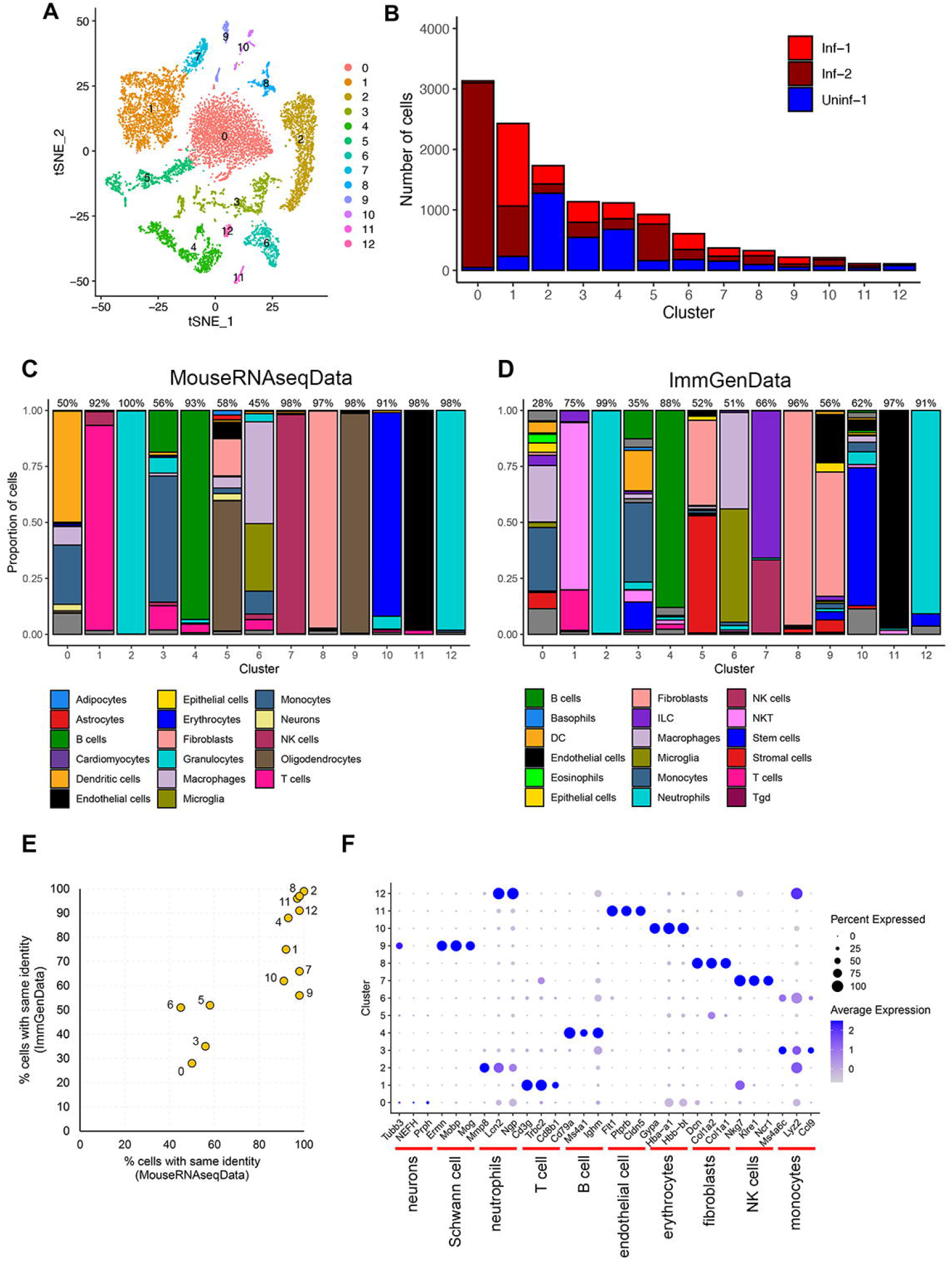
Clustering and annotation of cell populations. **(A)** Aggregated tSNE plot of all three datasets identifies 13 distinct clusters (0 – 12). **(B)** The proportion and total number of cells in each cluster shown differs between datasets e.g., cluster 0 is almost entirely composed of cells from the Inf-2 dataset. **(C and D)** SingleR was used to perform unbiased cell type recognition using both **(C)** MouseRNAseq and **(D)** lmmGen databases. For each cluster, the maximum proportion of cells given the same identity (analogous to a confidence score) is shown above the bar plot. **(E)** Both databases yielded similar results and confidence scores. Notably, clusters 0, 3, 5, and 6 could not be adequately resolved into a single dominant cell type. **(F)** Bubble plot showing both the proportion of cells in each cluster that express a specific marker and the expression level of that marker.

A second challenging component of scRNA-Seq analyses is the process of assigning specific cellular identities to a given cluster. This is typically achieved by identifying distinct markers within a given cluster and comparing this to a well-described reference database of cell identities. Here, we used the same annotation tool and reference databases as Wang *et al*. [8] (SingleR [25], with (MouseRNAseqData [26] and ImmGenData [27] from the celldex package (https://github.com/LTLA/celldex) (**Fig. 2C-D**). This analysis (i) produced convergent results (**Fig. 2E**) and (ii) demonstrated that while most clusters could be identified with high confidence (i.e., > 90% of cells present are predicted to have the same identity), other clusters are reported as a mix of cell types and thus should be considered as low confidence (i.e., Clusters 0, 3, 5, and 6, **Fig. 2C-D**). Subsequently, an analysis using representative markers for the cell types present in each cluster further confirmed the division between high confidence and low confidence cluster identities (**Fig. 2F**). We further analyzed Cluster 0 to better understand why no robust cellular identity could be assigned. Notably, this cluster derived almost entirely from the Inf-2 dataset (**Fig. 2B**) and, when compared to other clusters, was characterized as containing cells with high proportions of mitochondrial reads and low numbers of detectably expressed genes per cell (**Fig. 3A-B**). To test the hypothesis that most cells in this cluster are dying/dead, we summarized the expression of 40 cell death markers [28] and, again when compared to other clusters, determined these to be predominantly expressed in the Inf-2 derived cells in Cluster 0 (**Fig. 3C and Table S1**). This is particularly relevant in the context of the original Wang *et al*. [8] analyses as the major conclusions in that study were selectively derived from analysis of cells in this cluster.

**Figure 3.**
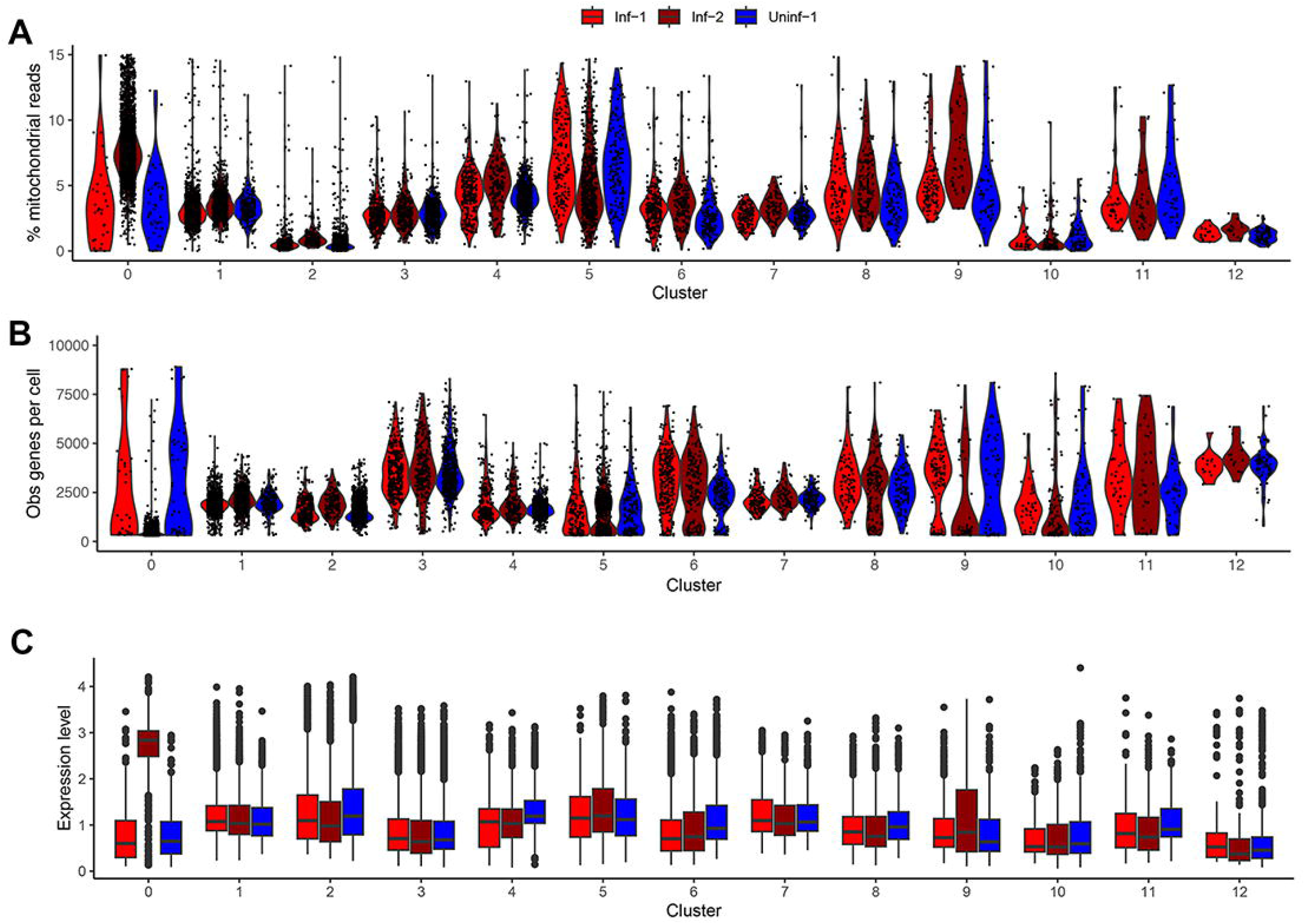
Further evaluation of cell quality within each cluster. For each cluster, and segregated by dataset, we determined **(A)** the proportion of mitochondrial reads per cell, **(B)** the number of distinct genes expressed per cell, and **(C)** the aggregated expression level of a selection of cell death markers (Table S1).

Taken together, these data indicate that (i) Inf-1 and Inf-2 datasets are not valid biological replicates, (ii) clusters 0, 3, 5, and 6 cannot be assigned a cell identity with confidence, and (iii) Inf-2 derived cells in Cluster 0 are likely undergoing programmed cell death.

### Off-target capture enables profiling of *LAT* intron lariats by scRNA-Seq

We next switched focus to the reported detection of HSV-1 *LAT* transcripts in multiple clusters. Of particular note is that the stable HSV-1 *LAT* 1.5 and 2 kb introns are not polyadenylated and the mature *LAT* RNA is highly unstable [15,18]. Thus, one would not expect to detect intron-derived RNAs by 3’ scRNA-Seq in which the 3’ oligo d(T) adapter is designed to prime of poly(A) tails. However, reanalysis of the raw fastq files from Wang *et al*. [8] demonstrated that 74 – 92 % of viral reads (representing < 0.005 % of all reads) aligned to the *LAT* intron, while the remaining reads mapped at low density throughout the rest of the HSV-1 genome (**Fig. 4A-B, Table S2)**. Closer examination of read alignments across the *LAT* locus showed consistent alignments that were associated with short adenosine homopolymers located within the intron and a much smaller peak at the 3’ end of the mature *LAT* (**Fig. 4C**). Taken together, these data show that off-target priming in 3’ scRNA-Seq experiments [29] surprisingly enables the detection of non-polyadenylated HSV-1 *LAT* introns. Similar results have been observed in other scRNA-Seq studies of HSV-1 latently infected ganglia, indicating that the 3’ scRNA-Seq approach is compatible with studies of HSV-1 latency models [7,30], however the efficiency of this off-target priming remains unknown.

**Figure 4.**
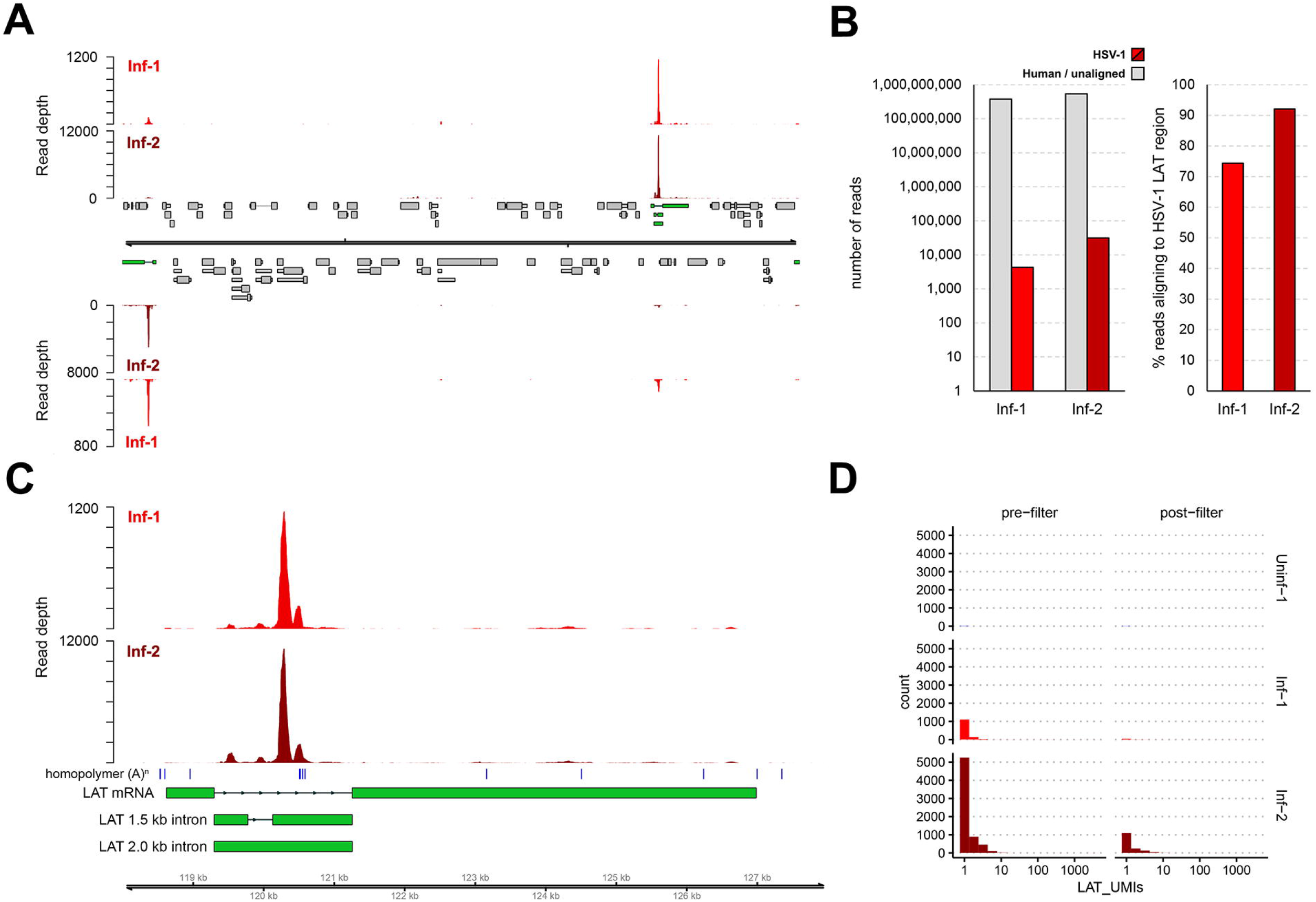
Abundant detection of non-polyadenylated *LAT* introns. **(A)** Coverage plot denoting the distribution of HSV-1 reads in the raw (unfiltered) Inf-1 and Inf-2 datasets. Double black lines represent the HSV-1 genome, wide and thin grey boxes indicate open reading frames and untranslated regions, and thin lines indicate introns. Top and bottom panel represent HSV-1 reads mapping to the forward and reverse strand of the genome. Both copies of the *LAT* locus are indicated in green. **(B)** Reads aligning to the HSV-1 genome comprised only a small proportion (< 0.006 %) of the Inf-1 and Inf-2 datasets, while most of these reads (74-92%) aligned to the *LAT* introns located in the *LAT* locus. **(C)** Coverage plot of the *LAT* locus confirms that the majority of HSV-1 *LAT* reads aligned next to short adenosine homopolymers (blue vertical bars) located within the intron, indicating off-target capture. (**D**) The majority of cells with *LAT* reads contained only a single copy of *LAT* (i.e., a single UMI) and most of these were removed during the QC filtering step.

### Loss of HSV-1 *LAT* during filtering suggests cell-free RNA contamination

*LAT* reads were not universally detected in the Inf-1 and Inf-2 datasets, but instead were 10 times more abundant in the Inf-2 dataset (**Fig. 4B**). In addition, most *LAT* reads in the raw dataset were excluded during the initial quality control filtering process (**Fig. 4D**). Notably, most cells with *LAT* reads in the filtered dataset only contained a single *LAT* read (as determined by the UMI present on each read) (**Fig. 4D**). Subsequent analysis of the individual clusters demonstrated that the vast majority of cells designated as *LAT+* were associated with Cluster 0 and were almost exclusively from the Inf-2 dataset (**Fig. 5A**). Similarly, the relative expression of *LAT* was highest in Cluster 0 (**Fig. 5B**). Because (i) this cluster is composed of dead/dying cells and (ii) HSV-1 *LAT* introns accumulate to high levels in neurons [31], we hypothesized that high background levels of cell-free RNA – originating from HSV-1 infected neurons that were damaged during tissue processing – could be the source of *LAT* reads in non-neuronal cells. To test this hypothesis, we compared the number of reads aligning to HSV-1 *LAT* and several cell type specific markers in the pre-(**Fig. 1A**) and post-filtered (**Fig. 1B**) datasets. Strikingly, this analysis demonstrated a significant loss of both HSV-1 *LAT* and neuronal markers during filtering that was not observed for any of the other major cell types present (**Fig. 5C & Fig. S1**). Thus, extensive death of (HSV-1 latently-infected) neurons during TG processing is the most likely source of ambient RNA contamination [32,33].

**Figure 5.**
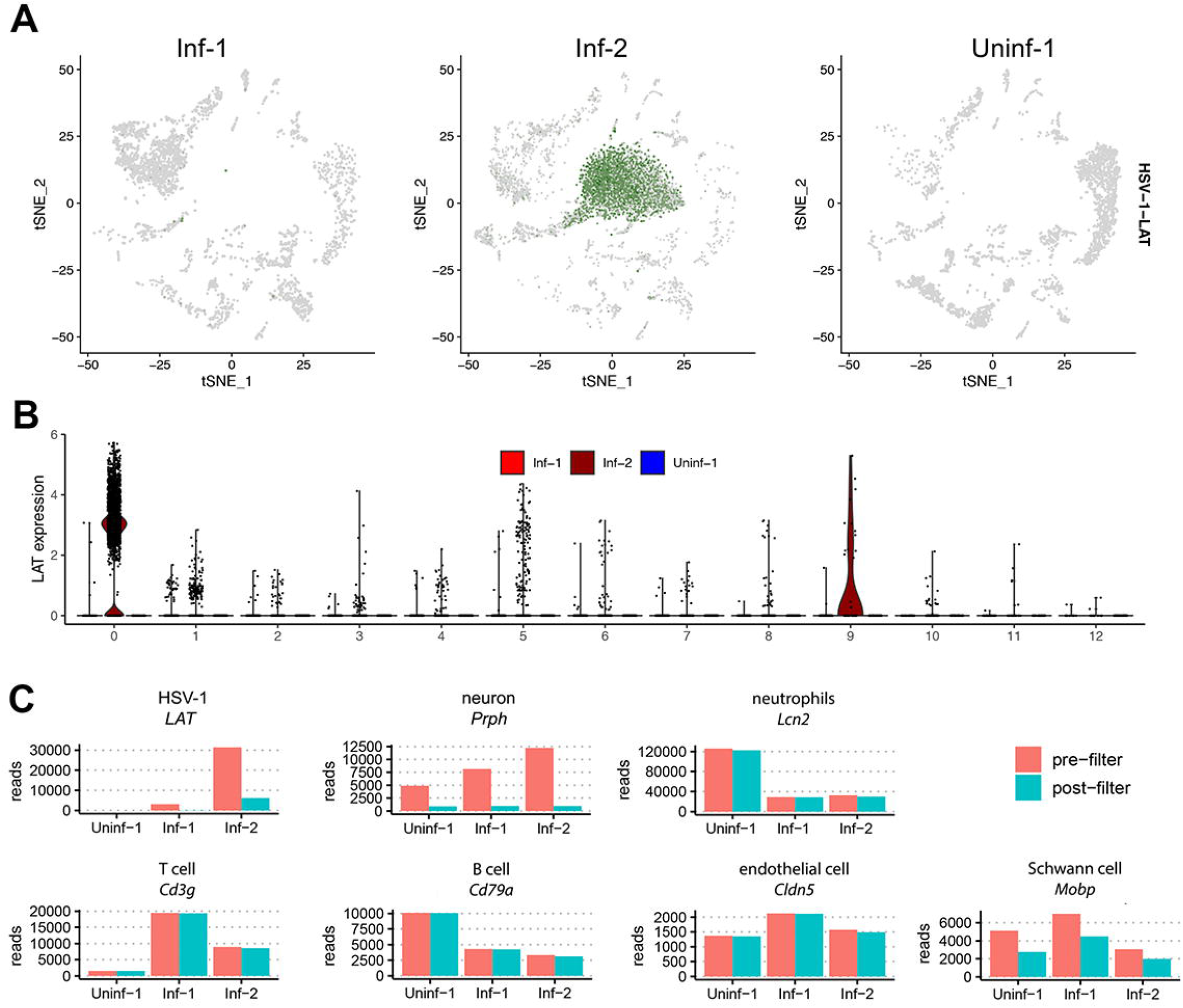
Abundant loss of reads associated with HSV-1 *LAT* and neuronal markers during filtering. (**A**) tSNE plot from Fig. 2A split by sample shows that the majority of *LAT* expression (in green) mapped to Cluster 0, which was exclusively present in the Inf-2 dataset. Violin plot showing log normalized expression of *LAT* in each cluster. (**C**) For HSV-1 *LAT* and representative markers of different cell types, we determined the total number of reads present in the pre-filtered (**Fig. 1A**) and post-filtered (**Fig. 1B**) datasets. Data for a wider selection of markers is shown in **Fig. S1**.

## Discussion

The recent study by Wang *et al*. [8] has challenged the dogma that HSV-1 exclusively establishes latency in neurons. However, overturning existing dogma necessitates robust and rigorous evidence that is supported by well-controlled independent experiments using orthologous methodologies. A key premise of Wang *et al*. [8] is that the presence of HSV-1 *LAT* RNA in a cell is sufficient to conclude that the virus has established latency in the infected cell. HSV-1 latency can be operationally defined as the presence of viral DNA in host cells in the absence of virus particle production, provided that the genome is maintained in a reactivatable state that enables the production of new infectious HSV-1 particles [34,35]. Transcriptional activity of the latent HSV-1 genome is repressed with exclusive expression of *LAT* driven by a neuron-specific promoter [11,13,15]. Neurons are the only cell type in which HSV-1 latency has been clinically and experimentally demonstrated in both human and mouse ganglia [2,9,36–39]. Interestingly, not all HSV-1 infected TG neurons express *LAT* in the HSV-1 mouse model [40] and it is unclear whether all TG neurons harboring HSV-1 DNA express *LAT* in humans [41,42], raising the question of whether all neurons containing HSV-1 DNA support virus reactivation. Thus, even if low-abundance HSV-1 *LAT* reads were detected in non-neuronal cells, this is not conclusive evidence of latency.

Our reanalysis of the scRNA-Seq data from Wang *et al*. [8] reveals significant errors in their data analysis. We conclude that their reported detection of *LAT* reads from non-neuronal cells is best explained by cell-free RNA originating from latently-infected neurons that were damaged during tissue processing. Moreover, we specifically establish that the Inf-1 and Inf-2 datasets cannot be considered biological replicates, with the Inf-2 dataset in particular showing evidence of extensive cell destruction during tissue processing. We have also shown that integration of these datasets yields multiple cell clusters that cannot be assigned a specific cellular identity. One of these clusters is dominated by low quality HSV-1 *LAT* expressing cells that are almost entirely derived from the Inf-2 dataset. A deep analysis of this cluster identified markers of multiple cell types including neurons and expression of a large number of programmed cell death markers. The significant loss of reads associated with HSV-1 *LAT* and neuronal markers during QC filtering further supports extensive neuronal cell death during tissue processing and the release of both neuronal RNAs and HSV-1 *LAT* into the homogenized single cell suspension. Such RNA is easily bound on the surface of other cell types and thus carried into the droplets in which cell lysis and the initial steps of scRNA-Seq library preparation take place [32,33].

As we had to specifically request that the scRNAseq datasets be made available to us post-publication, and these data are not linked to the publication on the journal website, we feel obligated to reiterate the FAIR (Findability, Accessibility, Interoperability and Reusability) data principles. These guidelines provide a framework to increase transparency and promote the reuse of data by the scientific community [43,44], which in turn will accelerate scientific discoveries. Many funding agencies, universities and scientific journals aim to promote open science by recommending or requiring researchers to adhere to the FAIR principles and open access publishing. In the context of scRNA-Seq experiments, this means that all raw sequence files, metadata, raw and filtered matrices, and all code/scripts used for analysis are deposited in publicly available data repositories (e.g., GEO, SRA, GitHub etc.), with references to the location of the data in the relevant sections of the article. It would have been helpful for Wang *et al*. [8] to include references to their scRNA datasets in the manuscript, following most journal guidelines, and to provide sufficient details in the Methods section to reproduce all aspects their data analysis e.g., the Cell Ranger parameters used for aligning raw sequence data and the construction of the hybrid genome reference were not described. While Methods sections are often written in a concise manner, it is increasingly common, and usually required by journals, that authors make available all scripts used for the analysis of the original data presented. Additionally, we recommend that authors demonstrate the impact of both QC filtering steps (such as those shown in **Fig. 1**) and cluster labelling strategies (**Fig. 3**) on each individual biological replicate. By showing this as Supporting Data in manuscripts describing scRNA-Seq data, it becomes easier for experts in the field and other interested parties to evaluate overall results.

In summary, our reanalysis of recently published scRNA-Seq data of HSV-infected mouse TG does not support the reported detection of HSV-1 *LAT* RNA in non-neuronal cells. While studies investigating the virus and host factors contributing to viral latency and reactivation at the single-cell resolution will undoubtedly advance our understanding of these processes, we encourage researchers to always adhere to the best practices for design and analysis of scRNA-Seq data, consider the biology of both the virus and host, and to share both the data and code used with the scientific community.

## Acknowledgements

WJDO and GMGMV are supported by the National Institute of Allergy and Infectious Diseases of the National Institutes of Health under Award Number R01-AI151290. Computational analyses were supported by Fred Hutch Scientific Computing (NIH ORIP grant S10OD028685). PR and KRJ are supported by the National Institute of Allergy and Infectious Diseases of the National Institutes of Health (R01AI132599). ALC is supported by an Australian NHMRC Level 3 Investigator grant no. APP1177942. DK and GMGMV are supported by NIH Contract 75N93019C00063. PRK acknowledges support of National Institute of Allergy and Infectious Diseases of the National Institutes of Health (R01-AI151290) and a National Eye Institute CORE Award (P30 EY08098). IM is supported by National Institutes of Health (NIH) grants R01-AI073898, R01-AI152543, and R01-GM056927. ACW is supported by the National Institute of Allergy and Infectious Diseases of the National Institutes of Health (R01-AI170583). DPD is supported by a German Centre for Infection Research (DZIF) Professorship and the National Institute of Allergy and Infectious Diseases of the National Institutes of Health (R01-AI170583, R01-AI152543).

## Methods

### Data sourcing

The raw data files (FASTQ) associated with the original study (SRA PRJNA937697) [8] were downloaded from the sequence read archive using fastq_dump from the SRA tools package (https://github.com/ncbi/sra-tools). Count matrices were downloaded from the Gene Expression Omnibus archive (GEO GSE225839) in order to reproduce the analyses in Wang *et al*. [8]

### Data processing and alignment

The 10x Genomics 3’ v3.1 datasets comprise three sets of reads, the I1 reads which contain the sample index, the R1 reads which contain the cellular barcodes and UMIs, and the R2 reads which contain the transcriptome sequences. To examine the nature and numbers of reads derived from the HSV-1 KOS transcriptome, we first performed quality and adapter trimming of the R2 reads using TrimGalore (--clip_R1 3 -q 30 --length 50) (https://github.com/FelixKrueger/TrimGalore) before aligning against the HSV-1 KOS genome (KT899744, [45]) using bbmap (https://sourceforge.net/projects/bbmap/). Resulting SAM files were parsed using SAMTools [46] and BEDTools [47] to generate bedgraph files that could be visualized in Rstudio using the packages GVIZ [48] and Genomic Features [49].

### Re-analysis of 10X data

Count matrices provided by the authors were imported and analyzed using the Seurat package in R [50]. Since the authors’ original analysis scripts were not provided, we attempted to use parameters from the manuscript where possible to recreate the analysis. However, Wang *et al*. [8] provided RNA- and UMI-level filter metrics for *LAT*-cells only, and it was unclear what filters were used on the full dataset. In our analysis, QC and filtering were performed on a combined dataset which included matrices from all three replicates (Uninf-1, Inf-1, and Inf-2). Mitochondrial filtering was performed at the same level as Wang *et al*. [8] (15%), and cells with 300-9000 features were included. Wang *et al*. [8]Scripts used in our reanalysis are available at https://github.com/proychou/10X_reanalysis. For tSNE and clustering we used parameters from Wang *et al*. [8] where available, e.g., top 20 principal components, resolution 0.1.

## Figure legends

**Figure S1.**
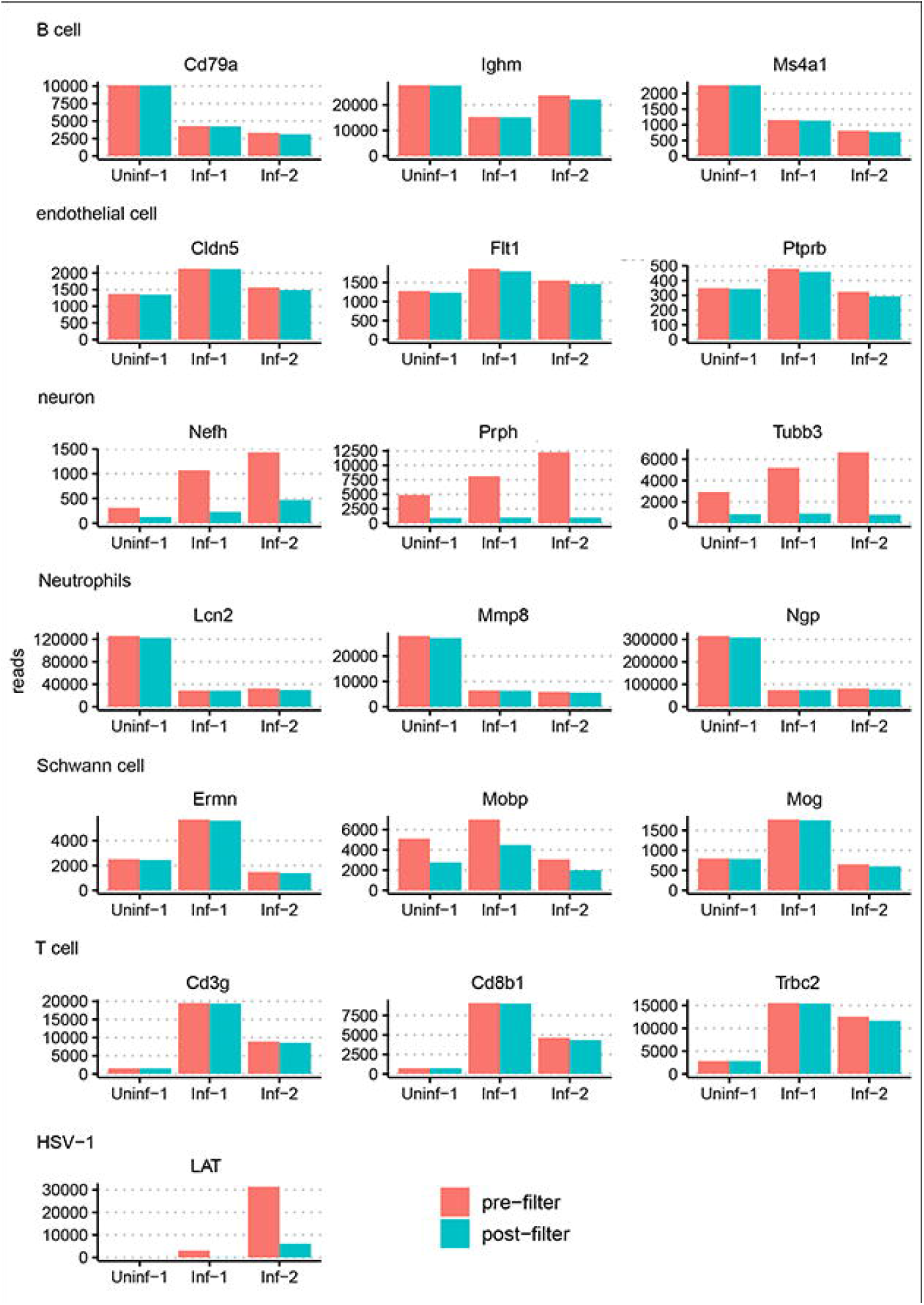
Further evidence that abundant loss of reads during filtering is specifically associated with HSV-1 *LAT* and neuronal markers. For HSV-1 *LAT* and representative markers of different cell types, we determined the total number of reads present in the pre-filtered and post-filtered datasets.

## References

1. Howley PM, Knipe DM, Mohr I, sodroski C. Herpes simplex viruses: mechanisms of lytic and latent infection. Fields Virology. Seventh Edition. Philadelphia: Lippincott Williams & Wilkins; p. 235–96.

2. Bastian FO, Rabson AS, Yee CL, Tralka TS. Herpesvirus hominis: isolation from human trigeminal ganglion. Science. 1972;178:306–7.

3. Vrabec JT, Payne DA. Prevalence of herpesviruses in cranial nerve ganglia. Acta Otolaryngol. 2001;121:831–5.

4. Nagel MA, Rempel A, Huntington J, Kim F, Choe A, Gilden D. Frequency and abundance of alphaherpesvirus DNA in human thoracic sympathetic ganglia. J Virol. 2014;88:8189–92.

5. Kennedy DP, Clement C, Arceneaux RL, Bhattacharjee PS, Huq TS, Hill JM. Ocular herpes simplex virus type 1: is the cornea a reservoir for viral latency or a fast pit stop? Cornea. 2011;30:251–9.

6. Remeijer L, Duan R, van Dun JM, Wefers Bettink MA, Osterhaus ADME, Verjans GMGM. Prevalence and clinical consequences of herpes simplex virus type 1 DNA in human cornea tissues. J Infect Dis. 2009;200:11–9.

7. Hu H-L, Srinivas KP, Wang S, Chao MV, Lionnet T, Mohr I, et al. Single-cell transcriptomics identifies Gadd45b as a regulator of herpesvirus-reactivating neurons. EMBO Rep. 2022;23:e53543.

8. Wang S, Song X, Rajewski A, Santiskulvong C, Ghiasi H. Stacking the odds: Multiple sites for HSV-1 latency. Sci Adv. 2023;9:eadf4904.

9. Stevens JG, Wagner EK, Devi-Rao GB, Cook ML, Feldman LT. RNA complementary to a herpesvirus alpha gene mRNA is prominent in latently infected neurons. Science. 1987;235:1056–9.

10. Rock DL, Nesburn AB, Ghiasi H, Ong J, Lewis TL, Lokensgard JR, et al. Detection of latency-related viral RNAs in trigeminal ganglia of rabbits latently infected with herpes simplex virus type 1. J Virol. 1987;61:3820–6.

11. Deatly AM, Spivack JG, Lavi E, Fraser NW. RNA from an immediate early region of the type 1 herpes simplex virus genome is present in the trigeminal ganglia of latently infected mice. Proc Natl Acad Sci U S A. 1987;84:3204–8.

12. Umbach JL, Nagel MA, Cohrs RJ, Gilden DH, Cullen BR. Analysis of human alphaherpesvirus microRNA expression in latently infected human trigeminal ganglia. J Virol. 2009;83:10677–83.

13. Depledge DP, Ouwendijk WJD, Sadaoka T, Braspenning SE, Mori Y, Cohrs RJ, et al. A spliced latency-associated VZV transcript maps antisense to the viral transactivator gene 61. Nat Commun. 2018;9:1167.

14. Umbach JL, Kramer MF, Jurak I, Karnowski HW, Coen DM, Cullen BR. MicroRNAs expressed by herpes simplex virus 1 during latent infection regulate viral mRNAs. Nature. 2008;454:780–3.

15. Zwaagstra JC, Ghiasi H, Slanina SM, Nesburn AB, Wheatley SC, Lillycrop K, et al. Activity of herpes simplex virus type 1 latency-associated transcript (LAT) promoter in neuron-derived cells: evidence for neuron specificity and for a large LAT transcript. J Virol. 1990;64:5019–28.

16. Puga A, Notkins AL. Continued expression of a poly(A)+ transcript of herpes simplex virus type 1 in trigeminal ganglia of latently infected mice. J Virol. 1987;61:1700–3.

17. Dobson AT, Sederati F, Devi-Rao G, Flanagan WM, Farrell MJ, Stevens JG, et al. Identification of the latency-associated transcript promoter by expression of rabbit beta-globin mRNA in mouse sensory nerve ganglia latently infected with a recombinant herpes simplex virus. J Virol. 1989;63:3844–51.

18. Devi-Rao GB, Goodart SA, Hecht LM, Rochford R, Rice MK, Wagner EK. Relationship between polyadenylated and nonpolyadenylated herpes simplex virus type 1 latencyassociated transcripts. J Virol. 1991;65:2179–90.

19. Wu TT, Su YH, Block TM, Taylor JM. Evidence that two latency-associated transcripts of herpes simplex virus type 1 are nonlinear. J Virol. 1996;70:5962–7.

20. Wu TT, Su YH, Block TM, Taylor JM. Atypical splicing of the latency-associated transcripts of herpes simplex type 1. Virology. 1998;243:140–9.

21. Phelan D, Barrozo ER, Bloom DC. HSV1 latent transcription and non-coding RNA: A critical retrospective. J Neuroimmunol. 2017;308:65–101.

22. Heumos L, Schaar AC, Lance C, Litinetskaya A, Drost F, Zappia L, et al. Best practices for single-cell analysis across modalities. Nat Rev Genet. 2023;1–23.

23. Ilicic T, Kim JK, Kolodziejczyk AA, Bagger FO, McCarthy DJ, Marioni JC, et al. Classification of low quality cells from single-cell RNA-seq data. Genome Biol. 2016;17:29.

24. Osorio D, Cai JJ. Systematic determination of the mitochondrial proportion in human and mice tissues for single-cell RNA-sequencing data quality control. Bioinformatics. 2021;37:963–7.

25. Aran D, Looney AP, Liu L, Wu E, Fong V, Hsu A, et al. Reference-based analysis of lung single-cell sequencing reveals a transitional profibrotic macrophage. Nat Immunol. 2019;20:163–72.

26. Benayoun BA, Pollina EA, Singh PP, Mahmoudi S, Harel I, Casey KM, et al. Remodeling of epigenome and transcriptome landscapes with aging in mice reveals widespread induction of inflammatory responses. Genome Res. 2019;29:697–709.

27. Heng TSP, Painter MW, Elpek K, Lukacs-Kornek V, Mauermann N, Turley SJ, et al. The Immunological Genome Project: networks of gene expression in immune cells. Nat Immunol. 2008;9:1091–4.

28. Díez J, Walter D, Muñoz-Pinedo C, Gabaldón T. DeathBase: a database on structure, evolution and function of proteins involved in apoptosis and other forms of cell death. Cell Death Differ. 2010;17:735–6.

29. Svoboda M, Frost HR, Bosco G. Internal oligo(dT) priming introduces systematic bias in bulk and single-cell RNA sequencing count data. NAR Genom Bioinform. 2022;4:qac035.

30. Aubert M, Strongin DE, Roychoudhury P, Loprieno MA, Haick AK, Klouser LM, et al. Gene editing and elimination of latent herpes simplex virus in vivo. Nat Commun. 2020;11:4148.

31. Kramer MF, Coen DM. Quantification of transcripts from the ICP4 and thymidine kinase genes in mouse ganglia latently infected with herpes simplex virus. J Virol. 1995;69:1389–99.

32. Young MD, Behjati S. SoupX removes ambient RNA contamination from droplet-based single-cell RNA sequencing data. Gigascience. 2020;9:giaa151.

33. Janssen P, Kliesmete Z, Vieth B, Adiconis X, Simmons S, Marshall J, et al. The effect of background noise and its removal on the analysis of single-cell expression data. Genome Biol. 2023;24:140.

34. Sawtell NM, Thompson RL. Herpes simplex virus and the lexicon of latency and reactivation: a call for defining terms and building an integrated collective framework. F1000Res. 2016;5:F1000 Faculty Rev-2038.

35. Cliffe AR, Wilson AC. Restarting Lytic Gene Transcription at the Onset of Herpes Simplex Virus Reactivation. J Virol. 2017;91:e01419–16.

36. Cook ML, Bastone VB, Stevens JG. Evidence that neurons harbor latent herpes simplex virus. Infect Immun. 1974;9:946–51.

37. Stevens JG, Cook ML. Latent herpes simplex virus in spinal ganglia of mice. Science. 1971;173:843–5.

38. Paine TF. Latent herpes simplex infection in man. Bacteriol Rev. 1964;28:472–9.

39. Goodpasture EW. Herpetic infection, with especial reference to involvement of the nervous system. 1929. Medicine (Baltimore). 1993;72:125–32; discussion 133-135.

40. Yang L, Voytek CC, Margolis TP. Immunohistochemical analysis of primary sensory neurons latently infected with herpes simplex virus type 1. J Virol. 2000;74:209–17.

41. Wang K, Lau TY, Morales M, Mont EK, Straus SE. Laser-capture microdissection: refining estimates of the quantity and distribution of latent herpes simplex virus 1 and varicella-zoster virus DNA in human trigeminal Ganglia at the single-cell level. J Virol. 2005;79:14079–87.

42. Maroui MA, Callé A, Cohen C, Streichenberger N, Texier P, Takissian J, et al. Latency Entry of Herpes Simplex Virus 1 Is Determined by the Interaction of Its Genome with the Nuclear Environment. PLoS Pathog. 2016;12:e1005834.

43. Wilkinson MD, Dumontier M, Aalbersberg IJJ, Appleton G, Axton M, Baak A, et al. The FAIR Guiding Principles for scientific data management and stewardship. Sci Data. 2016;3:160018.

44. Rocca-Serra P, Gu W, Ioannidis V, Abbassi-Daloii T, Capella-Gutierrez S, Chandramouliswaran I, et al. The FAIR Cookbook - the essential resource for and by FAIR doers. Sci Data. 2023;10:292.

45. Colgrove RC, Liu X, Griffiths A, Raja P, Deluca NA, Newman RM, et al. History and genomic sequence analysis of the herpes simplex virus 1 KOS and KOS1.1 sub-strains. Virology. 2016;487:215–21.

46. Li H, Handsaker B, Wysoker A, Fennell T, Ruan J, Homer N, et al. The Sequence Alignment/Map format and SAMtools. Bioinformatics. 2009;25:2078–9.

47. Quinlan AR, Hall IM. BEDTools: a flexible suite of utilities for comparing genomic features. Bioinformatics. 2010;26:841–2.

48. Hahne F, Ivanek R. Visualizing Genomic Data Using Gviz and Bioconductor. Methods Mol Biol. 2016;1418:335–51.

49. Lawrence M, Huber W, Pagès H, Aboyoun P, Carlson M, Gentleman R, et al. Software for computing and annotating genomic ranges. PLoS Comput Biol. 2013;9:e1003118.

50. Stuart T, Butler A, Hoffman P, Hafemeister C, Papalexi E, Mauck WM, et al. Comprehensive Integration of Single-Cell Data. Cell. 2019;177:1888–1902.e21.

